# Dynamic interhemispheric coordination in face processing

**DOI:** 10.1101/151076

**Authors:** Zhengang Lu, Bingbing Guo, Ming Meng

## Abstract

Our conscious experience of the world is normally unified. The brain coordinates different processes from the left and right hemispheres into one experience. However, the neural mechanisms underlying interhemispheric coordination remain poorly understood. A mechanistic approach to understanding interhemispheric coordination is “communication through coherence” (Fries, 2005; 2015). Using a recently developed time-resolved psychophysics (Fiebelkorn, Saalmann, & Kastner, 2013; Landau & Fries, 2012; Song, Meng, Chen, Zhou, & Luo, 2014), combined with fMRI decoding method, we investigated the interhemispheric coordination through coherence, by focusing on a quintessential case of hemispheric lateralized brain function: face processing in the left and right fusiform face area (FFA). We observed coherent oscillatory fMRI multi-voxel patterns in the left and right FFA when two stimuli presented successively cross visual fields, either initiating coordination from the left hemisphere or right hemisphere. When interhemispheric coordination started from the dominant right hemisphere, a coherent 44° phase difference between the left and right FFA in 3-4 Hz was observed; whereas when interhemispheric coordination started from the non-dominant left hemisphere, a coherent −17° phase difference between the left and right FFA in 5.5-6.5 Hz was observed. These results suggest that different phase coherence might mediate the interhemispheric coordination of face perception, depending on whether the initiating hemisphere is dominant or non-dominant. Our findings provide compelling fMRI evidence for interhemispheric coordination through coherence. The time-resolved fMRI decoding approach would be a useful starting point for a more promising approach for future investigation in interhemispheric dynamic coordination with fine-grained spatial and temporal resolution.

---

Our conscious visual experience is normally unitary despite response functions to stimuli such as words and faces are known to be different in the two cerebral hemispheres. Interhemispheric coordination is critical to accomplishing tasks that involve processing for which different hemispheres are specialized in a large range of behaviors and cognitive functions (Hellige, 1990). Moreover, split-brain patients had difficulties in combining information presented dividedly to the two hemispheres (Gazzaniga, 2000; Gazzaniga, Bogen, & Sperry, 1962), suggesting that interhemispheric coordination plays a vital role for normal unified conscious experience (Gazzaniga, 2005). Although the hemispheric specialization of human brain function has been demonstrated to exist for centuries, the neural mechanisms underlying how the two hemispheres *interact* to coordinate processes that are functionally-lateralized remain poorly understood. Investigating interhemispheric coordination is very important for understanding the what and how of human hemispheric specialization.

A powerful concept of understanding dynamic coordination in the brain is “communication through coherence (CTC)” (Fries, 2005; 2015). The theory of CTC suggests that dynamic relationships between remote brain areas can be modulated at fine timescales by the degree of coherence between oscillations in participating areas. This CTC scheme has been successfully used to study dynamic coordination with distant brain regions (Moser et al., 2010). Here, we extended the previous CTC studies to the interhemispheric spatial scale, that is, using CTC as a frame work to investigate the dynamic coordination between homotopic (corresponding) regions of the two hemispheres. A quintessential case of hemispheric specialization in brain function is that the left and right fusiform face areas (FFAs) play different roles in face perception (Bi, Chen, Zhou, He, & Fang, 2014; Kanwisher, McDermott, & Chun, 1997; McCarthy, Puce, Gore, & Allison, 1997; Meng, Cherian, Singal, & Sinha, 2012; Rossion et al., 2000; Sergent, 1985; Sergent, Ohta, & MacDonald, 1992). That hemispheric specialization has been observed at all stages of face processing (Rhodes, 1985) requires ongoing perceptual processing of face dynamically coordinated between the two hemispheres to give rise to a unified experience of face percept. Therefore, face processing in the right and left FFA provides an ideal candidate domain to reveal the neural mechanisms underlying interhemispheric dynamic coordination.

To investigate anatomically-precise interhemispheric dynamic coordination requires a measurement with both sufficient temporal and spatial resolution. fMRI measures BOLD signal simultaneously across the whole brain at the millimeter-level spatial resolution, therefore can be potentially used to examine dynamic interhemispheric coordination at a much larger spatial scale and higher spatial resolution than was previously possible. However, a temporal dynamic approach was previously hampered to applied to fMRI data due to the signal sampling limitations of fMRI. A newly developed time-resolved experimentation technique has been successfully used to assess the fine temporal profile in behavioral priming studies (Fiebelkorn et al., 2013; Huang, Chen, & Luo, 2015; Landau & Fries, 2012; Song et al., 2014). Here, we adopt this time-resolved experimental design with fMRI measurement of brain activities in both the left and right FFAs. The performance of a multivariate pattern classifier applied to fMRI responses was used to quantify the face information represented in the left and right FFA. This approach allowed the simultaneous recordings of the time-courses of brain activation in both the left and right hemispheres, tracking the interhemispheric relations in processing face information.

Twenty-two participants completed two or three fMRI sessions on separate days. Face/house localizer was used to functionally define the left and right FFA and other face-responsive brain areas in the left and right hemispheres. We expected that interhemispheric coordination initiated in the left hemisphere would be different from interhemispheric coordination initiated in the right hemisphere, due to the hemispheric specialization in face perception. In the main experiment, we used a divided field presented, cross visual field category matching task, in which the observers have to make a judgment on two images of face and/or house successively presented in the left or right side of periphery in the visual field (Gazzaniga & Smylie, 1983), whether they were the same category or not. Critically, the inter-stimulus-interval (ISI), or repetition lag, was varied from 0.033 to 1.333 s in steps of 0.033 s (Figure 1). The repetition effect, computed by comparing the multivariate classification accuracy for repeated faces condition with the classification accuracy for non-repeated face condition, was used to indicate the face information represented in the left and right face processing areas. These repetition effects were plotted separately as a function of repetition lags, and was transformed from the time domain to the frequency domain to further examine the coherence between the left and right face processing areas by analyzing the phase difference between the time courses of face representation in the left and right brain areas.

**Figure 1.**
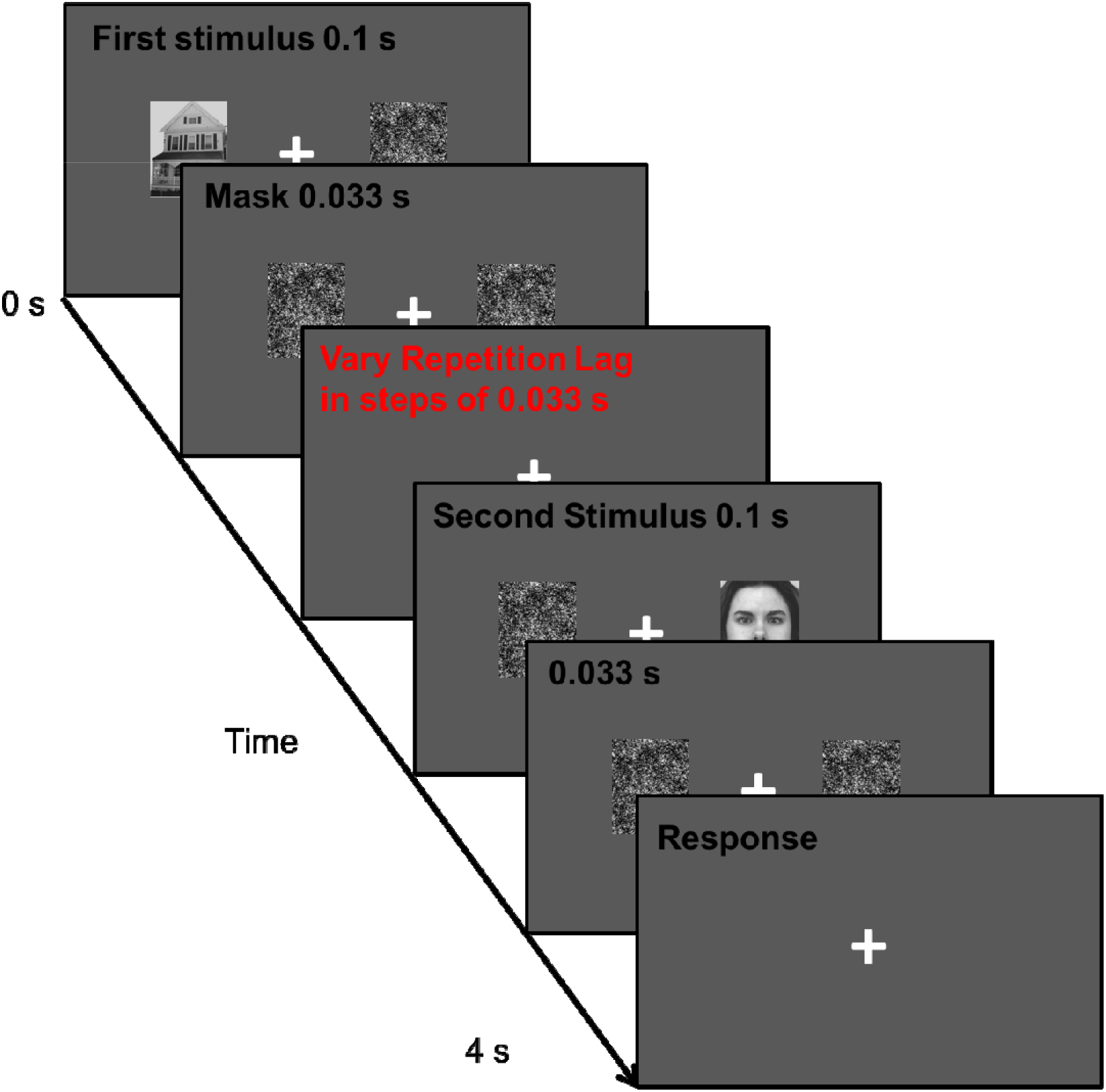
Experimental design. A typical trial was 4 s long. In each trail, the first display consisted a stimulus image (a face or a house) presented to the left (or right) visual field and a noise image side by side for 0.1 s. The stimulus and noise images were then backward masked by a pair of noise images presented side by side for 0.033 s. A second display consisted a stimulus image (a face or house) presented to the contralateral visual field with a noise image on the other side for 0.033 s, backward masked by a 0.033-s display of another pair of noise images. Critically, the first-to-second repetition lag ranged from 0.033 s to 1.333 s in steps of 0.033 s.

## Results

All observers maintained high performance accuracy for the category-matching task during scanning (mean = 97±1.5%). Reaction times (RT) were plotted as a function of repetition lags for each of the stimuli sequences (FF, HF, HH, and FH) in each of the sequential spatial locations (LVF-RVF and RVF-LVF). To estimate the RT time course, we averaged the RT within a 0.066 s time window, starting with the stimuli whose repetition lags were from 0.033 s to 0.099 s. We then moved this 0.066 s time window forward by 0.033 s and calculated the next average RT (0.066-0.132 s), repeating this procedure through the duration of repetition lags (0.033-1.33 s). Figure 2A shows the smoothed traces resulting from this procedure, averaged across all observers. A face repetition facilitation effect was evident and robust across repetition lags in the RT time courses in both spatial locations (LVF-RVF, RVF-LVF), as observers reliably responded faster in the face repetition condition than in the non-repetition condition (FF, HF, respectively). In contrast, the effect was weak and unreliable across repetition lags when comparing the raw RT time course in the house repetition condition with non-repetition condition (Figure 2B). Given the known functional lateralization of face processing, the stronger repetition effect when matching two sequentially and contralaterally presented faces than houses is consistent with the notion that the two hemispheres interact for face perception. Further fMRI decoding analysis focused on cross-hemispheres face repetition effect in the left and right FFAs.

**Figure 2.**
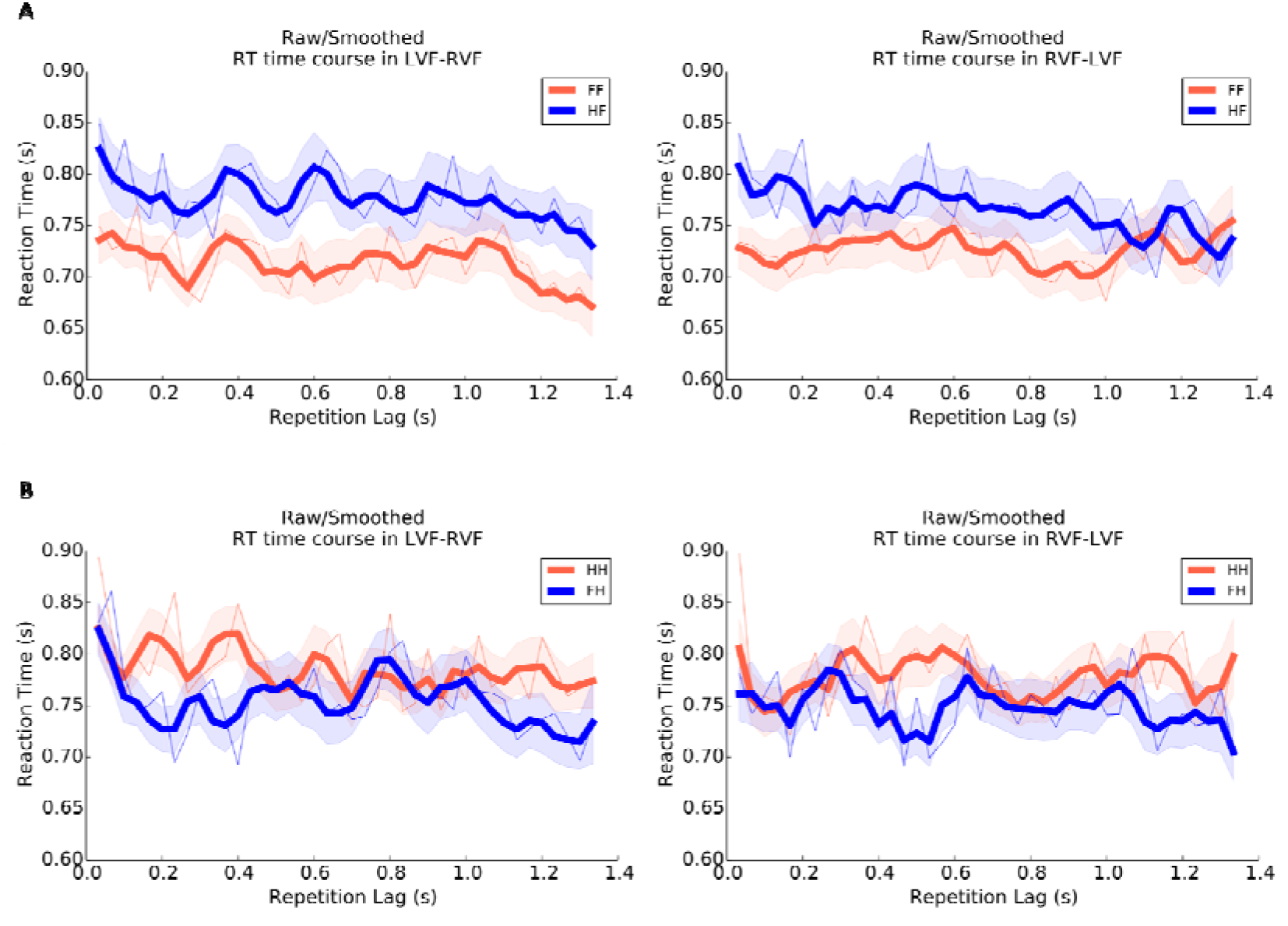
Behavioral RT results. **A**: RT results when the second stimulus was a face. Left: Average raw RT time courses as a function of first-to-second repetition lag (0.033-1.333 s in steps of 0.033 s) for face repetition (orange thin line) and non face repetition (blue thin line) conditions when the first stimulus presented to the left visual field, followed by the second stimulus presented to the right visual field. Average smoothed (0.066 s bin) RT time courses for face repetition (orange thick line) and non face repetition (blue thick line) conditions overlaid the raw RT time course. Right: Average raw and smoothed RT time course of face repetition and non face repetition conditions when the first stimulus presented to the right visual field, followed by the second stimulus presented to the left visual field. **B**: RT results when the second stimulus was a house. Left: Average raw RT time courses for house repetition (orange thin line) and non house repetition (blue thin line) conditions when the first stimulus presented to the left visual field, followed by the second stimulus presented to the right visual field. Average smoothed RT time courses for house repetition (orange thick line) and non house repetition (blue thick line) conditions overlaid the raw RT time course. Right: Average raw and smoothed RT time course of house repetition and non house repetition conditions when the first stimulus presented to the right visual field, followed by the second stimulus presented to the left visual field. Orange and blue shades indicate SEM.

The cross-hemispheres face repetition effects revealed by the facilitation in RTs suggests an interhemispheric coordination when stimuli were presented sequentially and contralaterally. To further reveal how the two hemispheres interact in face processing, fMRI response patterns were analyzed using multi-voxel pattern analysis (MVPA) in the left and right fusiform face areas (FFA) which are critically involved in face processing (Kanwisher et al., 1997). Activation patterns in the left and right PPA, which was reported to be involved in house processing (Epstein & Kanwisher, 1998)) were also analyzed for comparison. Since the goal of this study is to investigate the phase relationship between the left and right FFA, only data from observers who have both the left and right FFA were reported here (N = 20).

To estimate the time course of cross-hemisphere face repetition, the decoding accuracies as a function of repetition lags in different visual field sequences (LVF-RVF, RVF-LVF) were plotted for the left and right FFA in Figure 3. A clear face repetition effect was observed for the left and right FFA in both LVF-RVF and RVF-LVF, as the face repeated condition reliably revealed higher decoding accuracies than the non-repeated condition (except for the early stage of the time course). A cross-hemisphere face repetition index was formed in each ROI and each visual field condition by calculating the decoding accuracy difference between the face repetition condition and non-repetition condition (Figure 4). For each observer, a slowly developing trend was calculated by using 2nd order polynomial fit to the raw decoding accuracy difference time courses. Next, to remove possible interferences of this slow trend, and to better examine the oscillatory pattern, the slow trend was subtracted from corresponding decoding accuracy difference time courses for each observer to obtain detrended decoding accuracy difference time courses for each condition. Oscillatory patterns of face repetition effect in the left and right FFA under both LVF-RVF and RVF-LVF were evident in the resulting time course. To further measure the periodicity in the cross-hemisphere face repetition effect, the detrended time course data was converted into the frequency domain by using the fast Fourier transforms (FFT). A randomization procedure was performed to obtain the statistical thresholds by shuffling the time courses of face repetition effect independently for each observer 1000 times.

**Figure 3.**
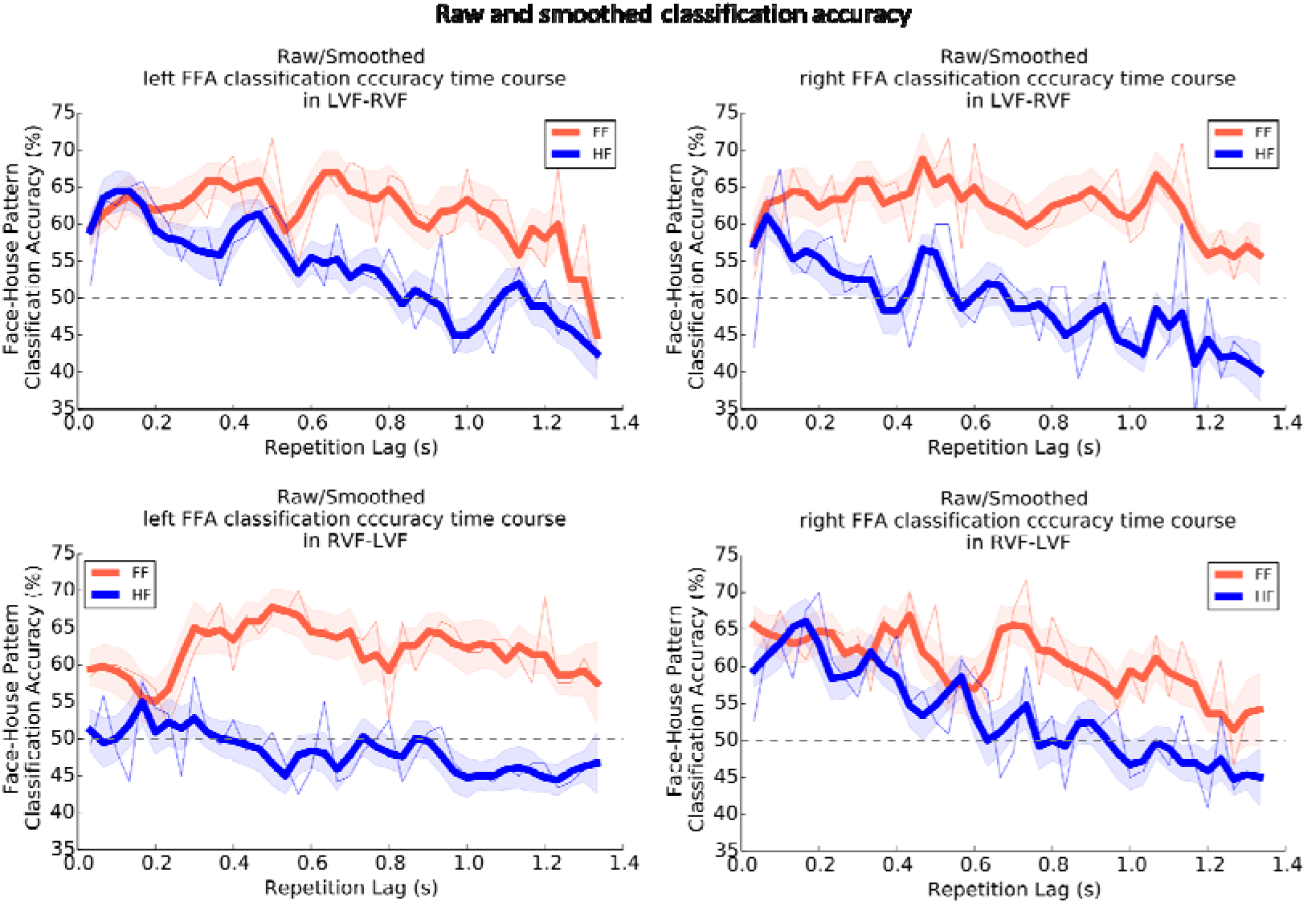
Oscillatory fMRI patterns in the FFA. Decoding results in the left and right FFA in both visual field conditions (LVF-RVF and RVF-LVF) for face repetition and non face repetition conditions. Each plot showed average raw classification accuracy time courses as a function of first-to-second repetition lag (0.033-1.333 s in step of 0.033 s) for face repetition (orange thin line) and non face repetition (blue thin line) conditions, overlaid by average smoothed (0.066 s bin) classification accuracy time courses for face repetition (orange thick line) and non face repetition (blue thick line) conditions. Gray dash lines indicate chance level. Orange and blue Shades indicates SEM.

**Figure 4.**
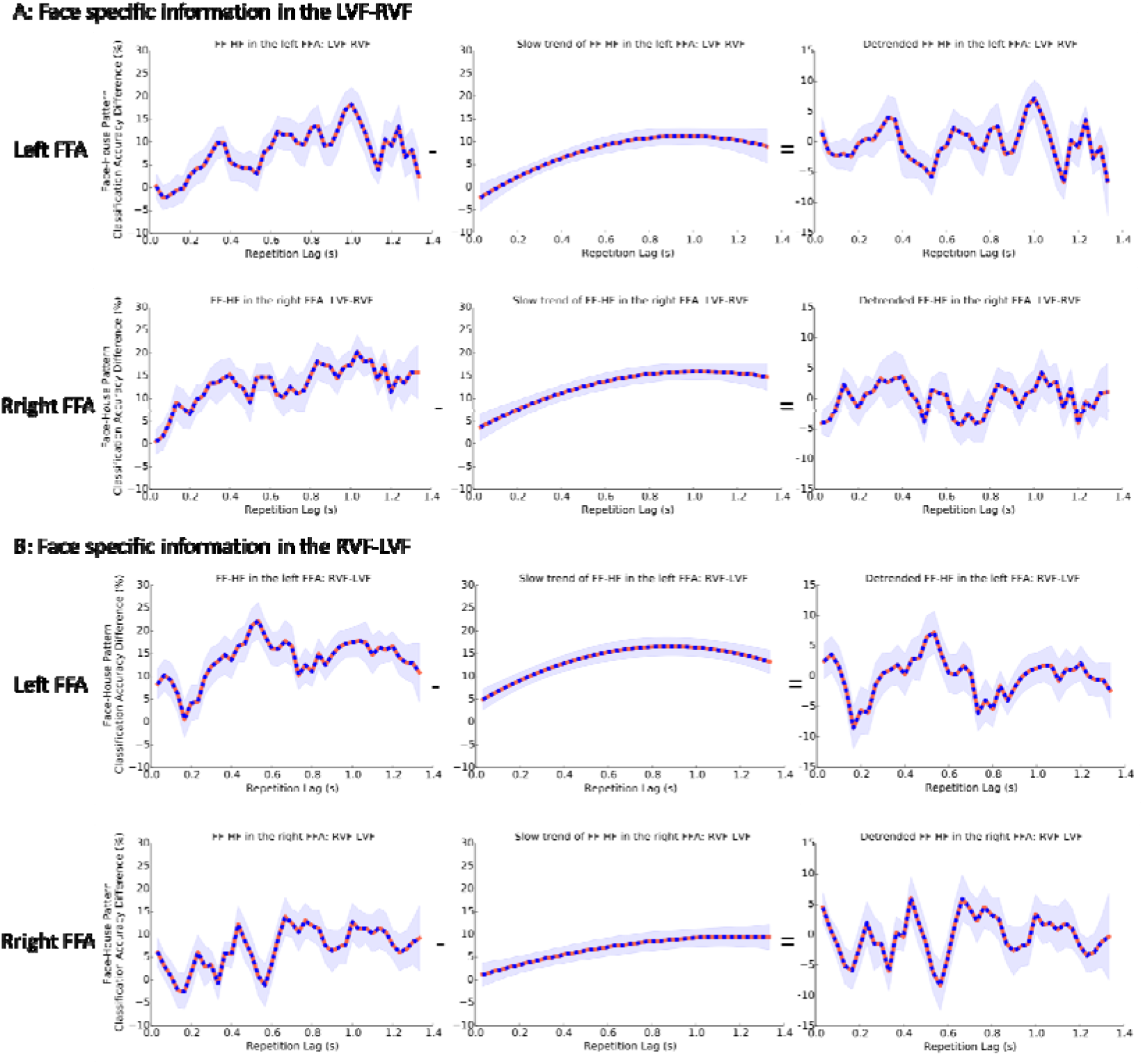
FF-HF decoding accuracy difference time courses. **A**: Face specific information represented in the left and right FFA in the LVF-RVF condition. Left: Face specific information time courses in the left and right FFA, calculated as the classification accuracy difference between face repetition and non face repletion conditions (FF-HF), as a function of first-to-second repetition lag (0.033-1.333 s in step of 0.033 s), smoothed with a 0.066 s bin. Middle: Slow trend averaged across all the observers. Right: Smoothed and detrended face specific information time courses in the left and right FFA, computed by subtracting slow trend (Middle) from smoothed time course (Left). **B**: Face specific information represented in the left and right FFA in the RVF-LVF condition. Blue shade indicates SEM.

In the LVF-RVF condition (Figure 5A and B), the left FFA revealed rhythmic fluctuation in face repetition effect in the theta-band, at approximately 5 Hz (permutation test, p < 0.05, for 4.9-5.8 Hz). In the right FFA, a similar rhythm was found (permutation test, p < 0.05, for 5.3-5.8 Hz). Importantly, the two rhythms had a significant phase relation that was clustered around 45° (Rayleigh test, p < 0.05, for 3-4 Hz), suggesting a coherent dynamic relation between the left and right FFA at theta-band when interhemispheric dynamics starting from the right hemisphere (Figure 6A). In the RVF-LVF condition, the spectral analysis revealed a significant amplitude peak in the right FFA at approximately 4 Hz (Figure 5B, permutation test, p < 0.05, for 3.9-4.7 Hz) and a significant peak in the left FFA at approximately 3 Hz (permutation test, p < 0.05, for 2.8-3.4 Hz). Interestingly, a significant phase relation, clustered around −17° (Rayleigh test, p < 0.05, for 5.5-6.5 Hz), was found between the left and right FFA when the interhemispheric dynamics starting from the left hemisphere (Figure 6B).

**Figure 5.**
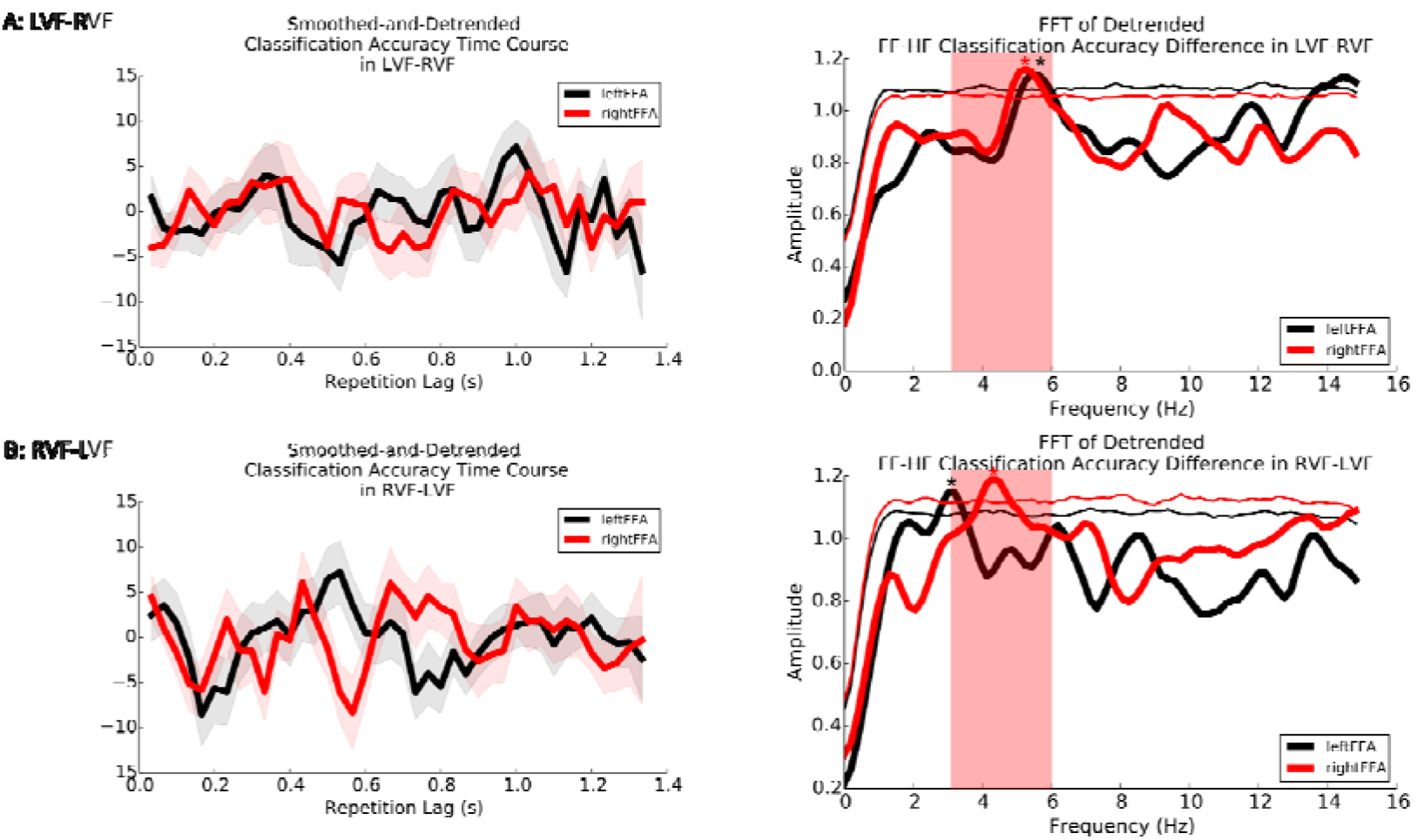
**A**: Oscillatory face specific information represented in the left and right FFA in the LVF-RVF condition. Left: Smoothed and detrended face specific information time courses in the left and right FFA (n = 20, mean ± SEM). Red and black shades indicate SEM. Right: Spectra for detrended time course of face-specific information (classification accuracy difference between FF and HF). Thin red and black lines indicate the statistical threshold of significance (*p* < 0.05, uncorrected), computed by performing a permutation test. Red rectangular shade indicates theta-range frequency (3-6 Hz). **B**: Oscillatory face specific information represented in the left and right FFA in the RVF-LVF condition. Smoothed and detrended time course of face-specific information shown in the left, and the frequency domain representation of the time course data shown in the right.

**Figure 6.**
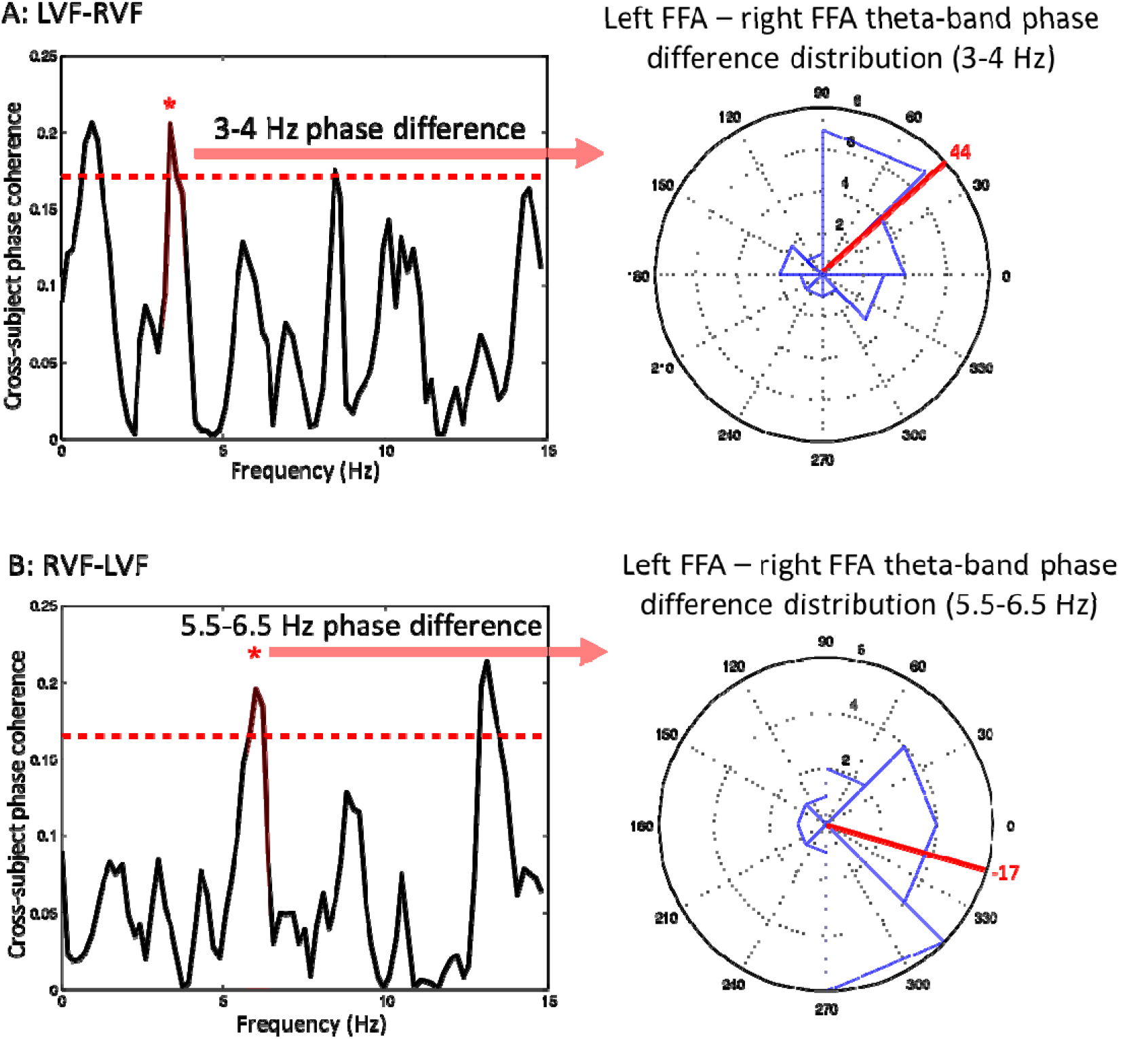
**A**: Left-Right FFA phase relationship in the LVF-RVF condition. Left: Cross-observers coherence in the phase difference between the left and right FFA as a function of frequency ranging from 0 – 15 Hz. The dash red line indicates the statistically significant threshold (*p* < 0.05) by performing permutation test and corrected for multiple comparisons. Right: Polar plot showing significant cross-hemisphere phase coherence at 3-4 Hz between the left and right FFA. The red line indicates significant 3-4 Hz phase difference between the left and right FFA averaged across observers (*p* < 0.05, Rayleigh test). **B**: Left-Right FFA phase relationship in the RVF-LVF condition. Left: Cross-observers coherence in the phase difference between the left and right FFA as a function of frequency ranging from 0 – 15 Hz. The dash red line indicates the statistically significant threshold (*p* < 0.05) by performing permutation test and corrected for multiple comparisons. Right: Polar plot showing significant cross-hemisphere phase coherence at 5.5-6.5 Hz between the left and right FFA. The red line indicates significant 5.5-6.5 Hz phase difference between the left and right FFA averaged across observers (*p* < 0.05, Rayleigh test).

The same FFT and phase difference analysis were performed on the fMRI pattern decoding accuracies in the PPA (Figure 7D). Instead of calculating the decoding performance difference between FF and HF in the PPA, the decoding accuracy difference was calculated between HH and FH, indicating the house repetition effect in the PPA. In contrast with the findings in the left and right FFA, no significant theta-band rhythmic fluctuation in house repetition effect was observed in neither the left nor right PPA in any visual field condition. Although significant alpha band fluctuation was observed in the right PPA under both visual field conditions, at approximately 10 Hz, the possibility of communication between the left and right PPA was excluded from further phase difference analysis, because no significant rhythmic fluctuation was observed in the left PPA in neither of the visual field conditions. Similarly, no significant simultaneously left and right theta-band rhythmic fluctuation of face repetition effects (decoding accuracy difference between FF and HF) was observed in the OFA, in the early visual areas, or in the entire temporal lobe in either of visual field conditions, therefore, no further phase difference analysis was performed (Figure 7A, B, and C).

**Figure 7.**
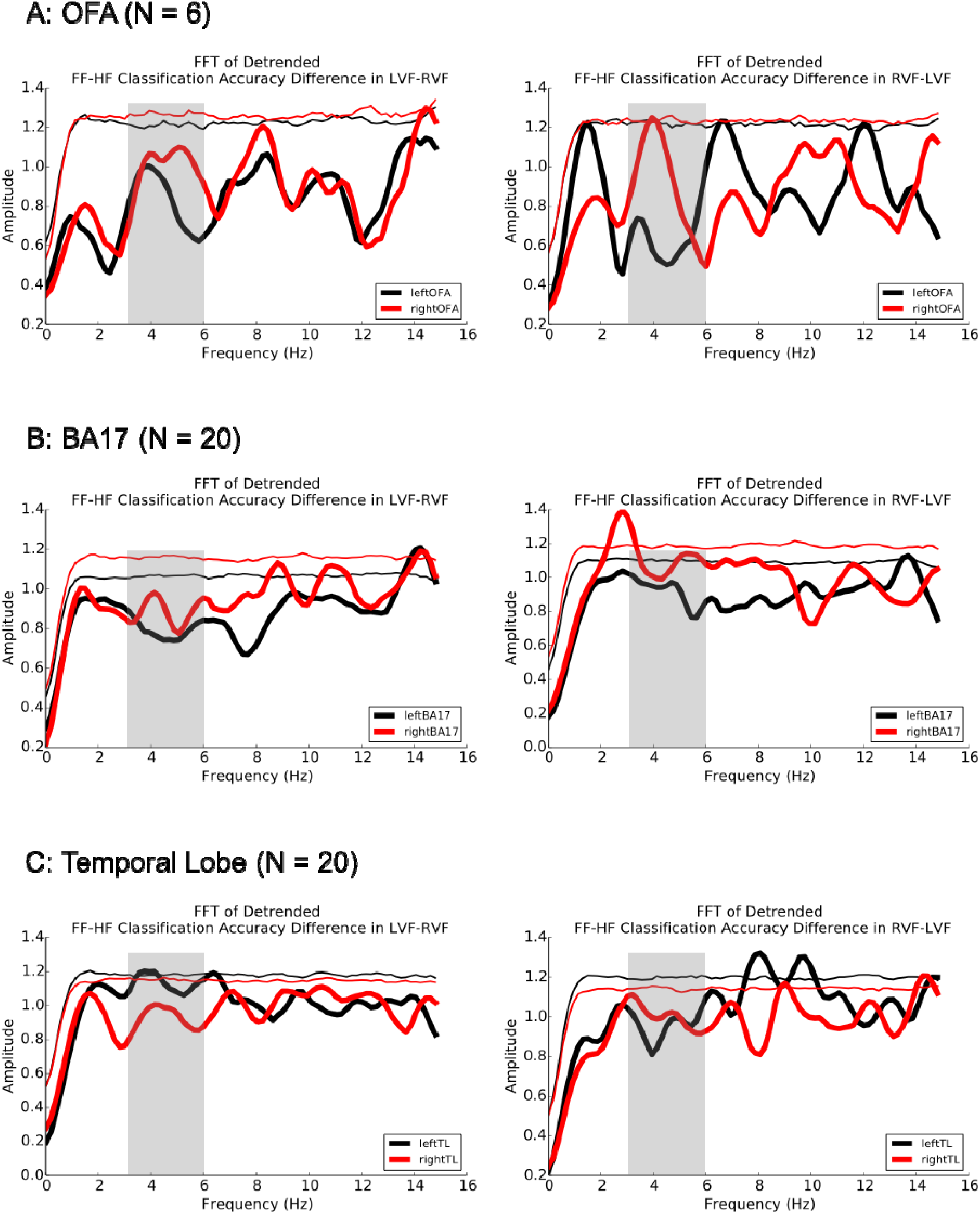
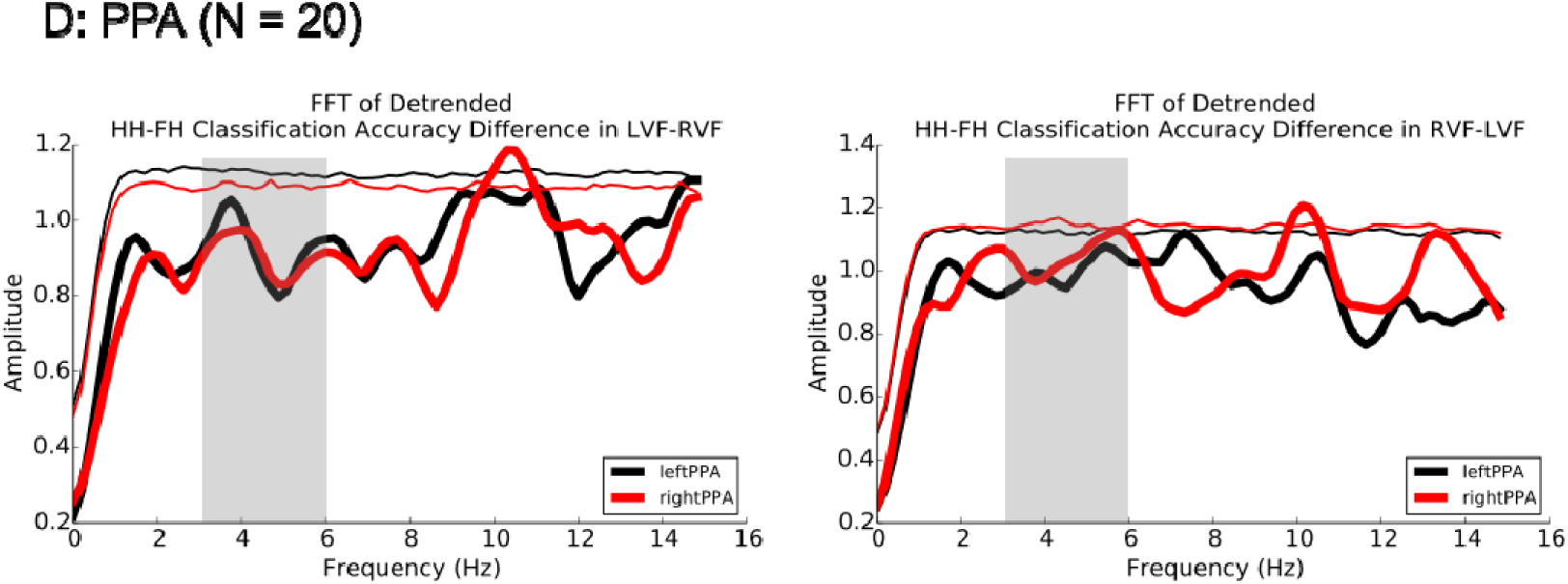
Frequency domain representation of the time course data in other ROIs. A, B, and C: Spectra for detrended time course of face-specific information (classification accuracy difference between FF and HF), in bilateral OFA, bilateral BA17 and bilateral Temporal lobe, respectively. Thin red and black lines indicate the statistical threshold of significance (*p* < 0.05, uncorrected), computed by performing a permutation test. D: Spectra for detrended time course of house-specific information (classification accuracy difference between HH and FH) in the left and right PPA. Thin red and black lines indicate the statistical threshold of significance (*p* < 0.05, uncorrected), computed by performing a permutation test. Gray rectangular shades indicate theta-range frequency (3-6 Hz).

## Discussion

That the left and right FFA play critical but different roles in face processing has been reported in previous neuroimaging studies (Meng et al., 2012; Rossion et al., 2000). It is still unknown how the left and right FFA interact with each other and how their coordination is flexibly modulated to form a unified face percept. In the present fMRI study, we investigated the neural underpinnings of interhemispheric dynamic coordination between the left and right FFA in a cross-hemispheres category matching task. Using a time-resolved fMRI decoding approach, we found that the decoding accuracies in the left and right FFA fluctuated in the theta-band and coherent phase difference between the left and right FFA was observed in this frequency range, suggesting that the interhemispheric coordination in face perception may occur in the theta-frequency range. We further observed that the starting hemisphere of interhemispheric coordination would modulate the phase relations between the left and right FFA.

Our results are not only consistent with previous findings showing that the phase of oscillatory brain activities in the theta-frequency range is involved in regulating the perception of visual information (Busch & VanRullen, 2010; Busch, Dubois, & VanRullen, 2009; Huang et al., 2015; Song et al., 2014; VanRullen, Busch, Drewes, & Dubois, 2011), but also show that the relations between the theta phases of multi-voxel pattern signals from the left and right FFAs would be indicative of the interhemispheric coordination in face processing. These theta phase relations between the homotopic are as across hemispheres could be further modulated by whether the dynamic coordination started from the dominant hemisphere or non-dominant hemisphere. These different temporal properties of interhemispheric coordination can be shown more pronouncing by band-pass filtering the original signal in the left and right FFA in the corresponding frequency range (Figure 8). When the dynamic coordination started from the nondominant hemisphere, i.e., left FFA, the left and right FFA exhibit rhythmic excitability fluctuations in the theta-frequency range, and a coherent phase lag (-17°) between the left FFA and right FFA was observed in the range of 5.5-6.5 Hz (Figure 6B), suggesting that the interhemispheric dynamic coordination occurs with a small time lag between the peaks (Figure 8B). In this viewing context, it might be beneficial if the information was synchronized from non-dominant to dominant hemisphere in a fast fashion. In comparison, when the dynamic coordination started from dominant hemisphere, i.e., the right FFA, the left and right FFA exhibit rhythmic excitability fluctuations in the theta-band frequency, but a larger phase lag (45°) between the left and right FFA was observed in the 3-4 Hz range within theta-band (Figure 6A), indicating that the left and right FFA may coordinate with a larger time lag between the peaks (Figure 8A). In this viewing context, it was likely to be beneficial for performing the matching task if the dominant hemisphere could coordinate with the non-dominant hemisphere, but the advantage of synchronizing the information was not as much as if the coordination started from a nondominant hemisphere.

**Figure 8.**
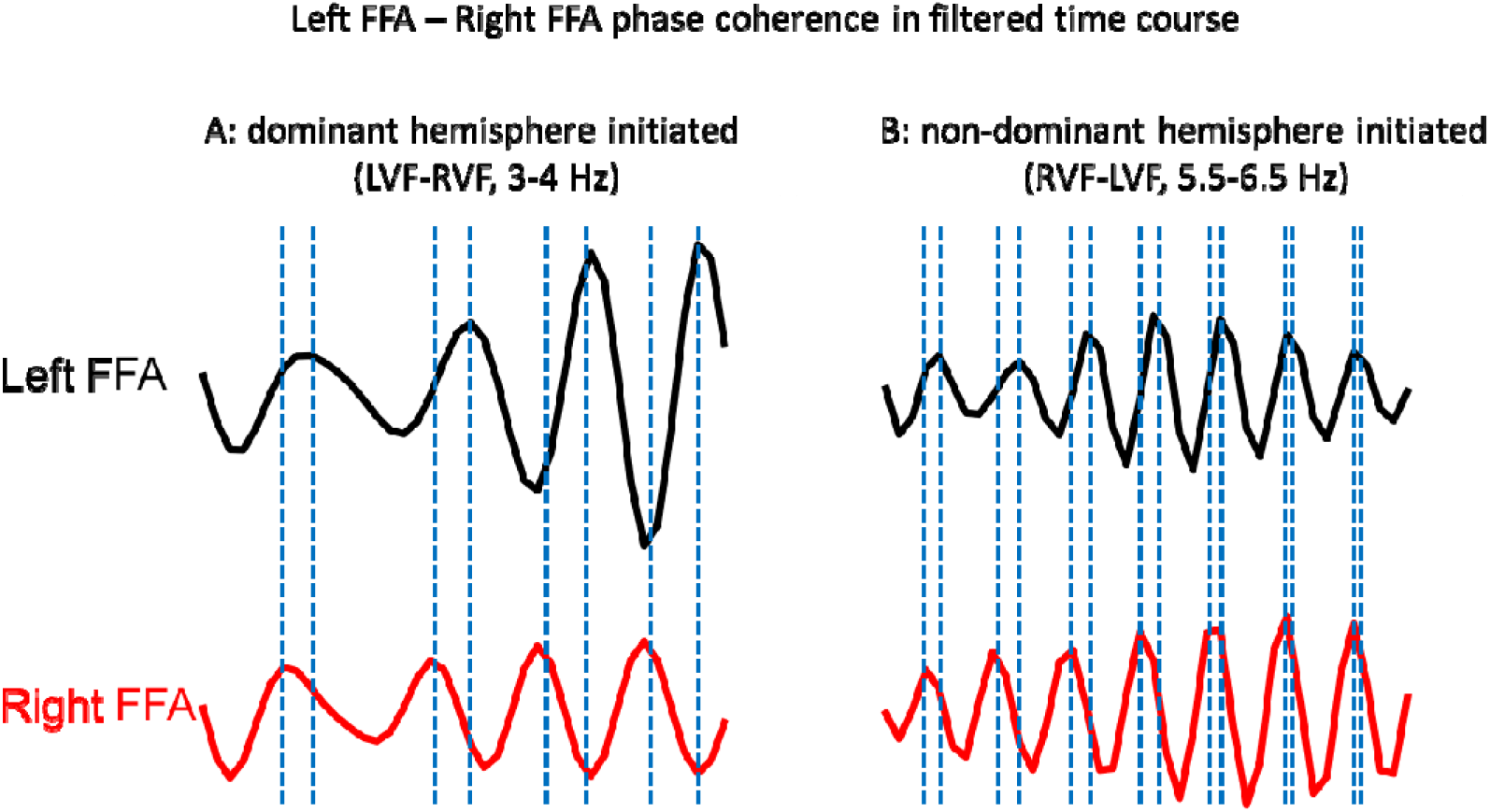
Frequency specific phase difference in fMRI pattern time courses. **A**: In the LVF-RVF condition (interhemispheric coordination initiated in the dominant hemisphere), the 3-4 Hz bandpass filtered and normalized FF-HF classification accuracy difference time course as a function of first-to-second repetition lag (0.033-1.333 s in step of 0.033 s) in the left and right FFA. **B**: In the RVF-LVF condition (interhemispheric coordination initiated in the non-dominant hemisphere), the 5.5-6.5 Hz band-pass filtered and normalized FF-HF classification accuracy difference time course as a function of first-to-second repetition lag (0.033-1.333 s in step of 0.033 s) in the left and right FFA. Blue dash lines indicate the coherent phase difference in corresponding specific frequency range.

The present study is the first to identify interhemispheric dynamic coordination by using fMRI measurement. Neuronal communication is subserved by neuronal coherence (“communication through coherence, CTC)” (Fries, 2005; 2015). When two brain areas oscillate coherently, the temporal windows are open for input and output from these two areas at the same time, resulting in an effective communication between the two areas (Engel, Gerloff, Hilgetag, & Nolte, 2013; Fries, 2005; Siegel, Buschman, & Miller, 2015). Coordinating interactions can occur among remote areas that generate coherent overall activity pattern (Moser et al., 2010). This CTC theory has been supported by recent studies on a variety of neural systems and rhythms using LFP, EEG and MEG (Catanese, Carmichael, & van der Meer, 2016; Fries, 2005; Siegel, Donner, & Engel, 2012). An limitation in these study is that, these methods are prone to be confounded by signal mixing artifacts due to their limited spatial resolution (Engel et al., 2013; Nolte et al., 2004; Stam, Nolte, & Daffertshofer, 2007), which is especially severe for measuring anatomically-precise brain interactions, for example, the dynamic interaction between left and right FFA. Our study provided a feasible approach to investigate dynamic coordination among anatomically-precise brain regions cross hemispheres with high temporal resolution. First, to achieve a dense temporal assessment of neural activity in the left and right face areas when interhemispheric coordination occurred, we combined a cross-hemispheres fMRI repetition paradigm (Grill-Spector & Malach, 2001; Grill-Spector, Henson, & Ca, 2006) with a recently developed time-resolved psychophysical approach, which has successfully probed the fine temporal dynamics in behavioral performance (Fiebelkorn et al., 2013; Huang et al., 2015; Landau & Fries, 2012; Song et al., 2014). Second, since neural activity pattern have better temporal resolution than univariate BOLD signal change (Kohler et al., 2013), to further enhance the sensitivity of the time-resolved fMRI design, we applied multi-voxel pattern decoding technique to the fMRI data, and analyzed the time course of decoding accuracies to reveal the interhemispheric dynamic coordination. Taken together, the time-resolved fMRI decoding approach used in this study successfully captured the fine spatio-temporal dynamic of pattern fluctuations in the left and right FFA, and revealed the dynamic coordination between these two regions.

To conclude, our results provide a space- and time-resolved view of interhemispheric coordination in face perception. We showed that the interhemispheric dynamic coordination between the left and right FFA may occur at theta-frequency range, and the dynamic coordination between the two are modulated by whether or not the initiating hemisphere is dominant in face processing. The time-resolved fMRI decoding approach provides a useful tool to investigate the spatio-temporal dynamics in the brain systems.

## Methods

### Observers

Twenty-two observers (18 right-handed; 14 females; mean age = 21.6 ± 3.6 years) participated in the experiment. All observers had normal or corrected-to-normal visual acuity. They were recruited from Dartmouth College community, gave written informed consent before participation, and received course credits or monetary payment for their participation. The experiment was approved by the Dartmouth College Committee for the Protection of Human Subjects.

In addition to these participants, 5 observers were tested but excluded from further data analysis, 4 of them due to excessive head motion during the experiment and 1 due to the observer’s failure to maintain above chance level behavioral performance.

### Experimental Procedures

In the main experiment, a female face photograph and a frontal-view house photograph were used as stimuli, subtended 8.5 × 6.9° visual angle. Figure 2 shows the experimental design and stimuli presentation procedure. The experimental design was adapted from a fast, event-related fMRI repetition paradigm (Grill-Spector et al., 2006; Grill-Spector & Malach, 2001). In each trial, presented in a semi-randomized order (stimuli conditions counter-balanced between observers), observers viewed a sequential but contralateral presentation of two photographs, resulting in 2 spatial location sequences (left-to-right visual field: LVF-RVF, and right-to-left visual field: RVF-LVF) and 4 stimuli presentation sequences (face-face: FF, face-house: FH, house-face: HF, and house-house: HH). Moreover, similar to the study by (Towler & Eimer, 2015), in each display, a photograph and a noise image were presented simultaneously to the left and right of fixation (0.75° × 0.75°) at a horizontal eccentricity of approximately 6°, relative to the center of the photograph and the noise image. Each face (or house) photograph was presented for 0.1 s, and immediately followed by a 0.033-s noise image mask. Most critically, the inter-stimulus-interval (ISI), or repetition lag, was varied from 0.033 to 1.333 s in steps of 0.033 s, corresponding to a sampling frequency of 30 Hz. Each trial was four-second long with no inter-trial-interval. Within this time-window, observers were instructed to perform a modified one-back, same-different category task by using their left hand to make a “same” or “different” button-press while maintain fixating at the center of the screen. Each observer completed 1920 trials in total: 6 repetitions for each of the 4 stimuli presentation sequences in each of the 2 spatial location sequences at each of the 40 ISIs (from 0.033 to 1.330 s in step of 0.033 s). Each experimental run lasted 5 min 28 s, consisting of 80 trials with two 4-s fixation periods at the beginning and the end. Each observer completed 24 experimental runs in two or three sessions on separate days (4 observers completed in three sessions).

To localize brain regions of interest (ROI) that are involved in face processing, in separate scan runs, we presented observers with blocks of grayscale face and house photographs, subtended 8.5 × 6.9° visual angle. These face and house photographs never appeared in the main experiment. There were 10 sixteen-second-long stimulus blocks within a run, including 5 face blocks and 5 house blocks interleaved with 16 s fixation-only periods. The fixation-only periods were also included at the beginning and the end of each run. The presentation order of stimulus blocks was randomized for each observer. Each stimulus block contained 16 faces (or houses), each presented for 0.5 s followed by a 0.5 s fixation-only interval. To ensure attention to the displays, observers were asked to maintain fixation at the center of the screen and use their left hand to make a button-press whenever they detected a repeat of the stimulus in succession. Observers completed 4-6 ROI localizer runs in total, with each run lasting 5 min and 36 s. Two observers completed 5-6 runs in three sessions because their FFAs could not be localized reliably with 4 runs. The additional runs allowed us to obtain comparable size of the FFAs across all observers (see Table 1).

**Table 1.**
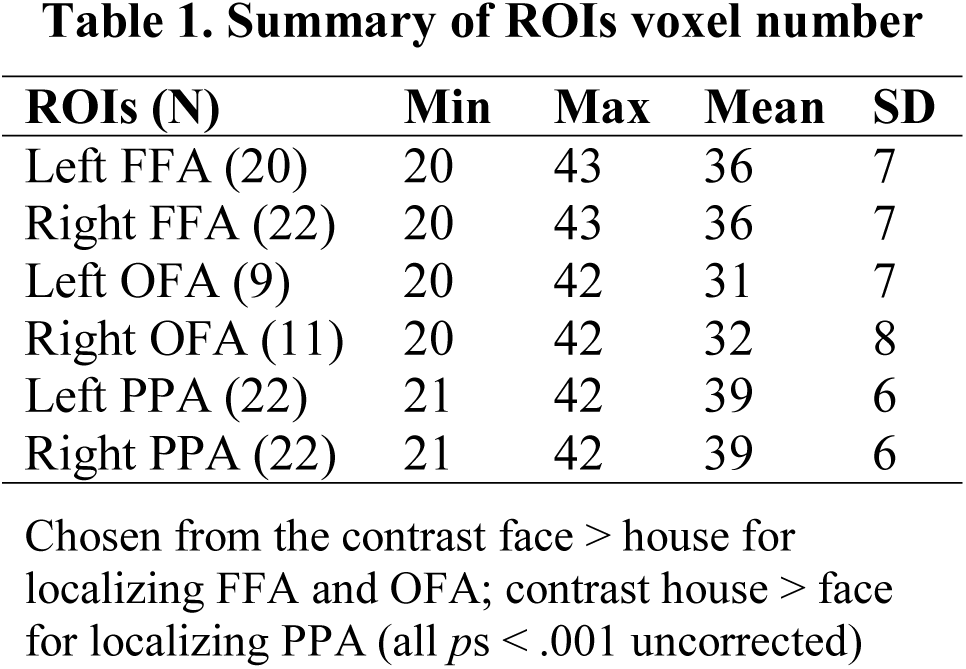
Summary of ROIs voxel number.

| ROIs (N) | Min | Max | Mean | SD |
| --- | --- | --- | --- | --- |
| Left FFA (20) | 20 | 43 | 36 | 7 |
| Right FFA (22) | 20 | 43 | 36 | 7 |
| Left OFA (9) | 20 | 42 | 31 | 7 |
| Right OFA (11) | 20 | 42 | 32 | 8 |
| Left PPA (22) | 21 | 42 | 39 | 6 |
| Right PPA (22) | 21 | 42 | 39 | 6 |

Chosen from the contrast face > house for localizing FFA and OFA; contrast house > face for localizing PPA (all $ps < .001$ uncorrected)

### fMRI Methods

fMRI data were acquired from a 3.0 T Philips Intera Achieva scanner (Philips Medical Systems, Bothell, WA) using a 32 channels head coil at the Dartmouth Brain Imaging Center (Hanover, NH). Stimuli were presented to observers via a Panasonic DT-4000U DLP projector. Observers viewed stimuli back-projected onto a screen at the rear of the scanner bore through a mirror mounted to the head coil. The width and height of the projected screen were 45.7 cm and 34.3 cm (1024 × 768 pixels) respectively. The distance between the mirror and the projected screen was 97.8 cm. The distance between observers’ eyes and mirror was approximately 12.7 cm. Stimuli presentation and the collection of behavioral responses were controlled by a IBM ThinkPad running MATLAB 2011b with Psychtoolbox (Brainard, 1997). Before functional imaging in each session, a high-resolution T1-weighted 3D-MPRAGE anatomical scan was acquired for each observer (FOV = 240 mm, TR = 8.2 ms, TE = 3.8 ms, flip angel = 8°, voxel size = 1 mm × 1 mm × 1mm, reconstruction matrix = 256 × 256, 222 slices) to allow the functional scans in each session to be registered to the observer’s high-resolution anatomical scan. Functional data were acquired using standard gradient-echo echo-planar imaging (EPI) sequence (FOV = 240 mm, TR = 2000 ms, TE = 35 ms, flip angel = 90°). Each volume of the fMRI data contained 35 slices (slice thickness = 3 mm, slice gap = 0.3 mm, in-plane resolution = 3 mm × 3 mm) tilted to ensure full coverage of occipital and temporal lobes. All functional images were acquired in an interleaved slice order.

### Data Analyses

Functional data were analyzed with AFNI (Cox, 1996) and in-house Python codes. Data preprocessing included transient spikes suppression, slice timing correction, head motion correction, spatial smoothing (4 mm full width at half maximum smoothing kernel), intensity normalization, linear drift correction, and Talairach space transformation (Talairach & Tournoux, 1998). To establish the voxel correspondence across sessions for each observer, data from two or three sessions were spatially aligned by applying the rotation parameters computed from aligning the anatomical scan of each session to the mean anatomical image. All data was analyzed in both the native space of each observer and Talairach space.

### ROI definitions

A whole-brain general linear model (GLM) analysis was performed separately on the fMRI data acquired from each observer in the ROI localizer scan runs. The fusiform face area (FFA) was defined as continuous clusters of 20 or more significant voxels in the fusiform gyrus showing greater activation for faces than for houses (*p* < 0.001, uncorrected) (Kanwisher et al., 1997). We were able to localize both the right and left FFAs in 20 out of the 22 observers (the left FFA was not found in 2 observers). Using the same contrast, we were also able to localize the occipital face area (OFA, (Gauthier et al., 2000; Puce, Allison, Asgari, Gore, & McCarthy, 1996), defined as continuous collection of 20 or more significant voxels in the occipital lobe in six observers with bilateral OFA, three observers with left OFA only, and five observers with right OFA only (*p* < 0.001, uncorrected). Following previously established procedures (Epstein & Kanwisher, 1998), the parahippocampal place area (PPA) was defined in all the observers as the collection of more than 20 continuous voxels in the bilateral collateral sulcus and parahippocampal gyrus showing greater activation for houses than for faces (*p* < 0.001, uncorrected). To control for any potential confounding effect of ROI size, the size of each ROI roughly matched (see details in Table 1).

In addition to the above-described ROIs, we created anatomically defined ROIs corresponding to the early visual areas and the entire temporal lobe (bilateral Brodmann Area 17: left BA17 was 95 voxels, right BA17 was 88 voxels; TT_N27 template: left temporal lobe was 200 voxels and right temporal lobe was 210 voxels). These additional ROIs were localized using AFNI’s atlas segmentation.

### Univariate analysis of averaged BOLD

The univariate analysis was used to determine the peak TR of fMRI responses to the stimuli. Data from this particular TR were then used to extract multivariate activation patterns in further analysis. Following the standard ROI-based analysis approach, ROI masks were overlaid onto the data from the main experiment and time courses from each observer were extracted by averaging the fMRI data over all the voxels in each of the defined ROIs. As in previous studies (Kourtzi & Kanwisher, 2001; Lu, Li, & Meng, 2016), the averaged time courses were converted to percent signal change by subtracting and dividing by a baseline value, which is the average of fMRI activity at 1 TR before the stimulus onset and the activity at the stimulus onset TR, and multiplying it by 100. We then collapsed the time courses of the visual field conditions and all the ISI conditions, and determined the time point of greatest signal amplitude in the averaged response. Consistent with previous reports, fMRI responses peaked at ~6 s after trial onset (Aguirre, Zarahn, & D’Esposito, 1998; Frahm, Bruhn, Merboldt, & Hanicke, 1992; Kwong et al., 1992; Miezin, Maccotta, Ollinger, Petersen, & Buckner, 2000). Multi-voxel activation patterns at this time point for each observer and every experimental trial were extracted from each ROI, and were then used in subsequent multivariate analyses.

### Multivariate pattern analysis (MVPA)

MVPA was performed using PyMVPA (Hanke et al., 2009). Decoding analysis was conducted on the delay period preprocessed fMRI response (6 s after trial onset) including each voxel in a given ROI, similar to a previous study (Lu et al., 2016). This time point was selected because it accounted for the hemodynamic response lag. fMRI responses across all the voxels in each ROI were first normalized using z-score transformation to scale all voxels into the same average in order to remove any effects related to overall amplitude differences, and then detrended to remove polynomial trends from the data. The resulting fMRI response pattern from each ROI was used in the decoding analysis. Using a leave-one-trial-out cross validation procedure, we trained a linear support vector machine (SVM) classifier to discriminate the category of the second stimulus (i.e., face or house) based on the response patterns from N-1 of the total N trials, and then the classifier assigned the response pattern from the Nth trial to one of the two stimulus category (i.e., face or house). This procedure was repeated N times with each of the N trials serving as testing pattern and the remaining N-1 trials as training patterns. Decoding accuracy was defined as the proportion of test patterns that were correctly classified into either face or house category, with 50% correct being chance level performance. Decoding accuracies were then averaged across all the training-testing procedures for each observer for subsequent spectrum frequency analyses.

### Analysis of frequency

Frequency analysis was performed with custom-written Python and MATLAB EEGLAB toolbox. In each observer, the temporal profile of decoding accuracies was derived as a function of repetition lag from 0.033 s to 1.33 s in steps of 0.033 s (30 Hz sampling frequency) for each experimental condition in each ROI. In order to extract the slow developing trend, raw averaged decoding accuracies for each experimental condition was fitted to a second order polynomial function. This slow trend was then subtracted from the corresponding temporal profile to obtain the detrended decoding accuracies of each experimental condition for each observer. A fast Fourier transformation (FFT) was performed to convert the temporal profile of detrended decoding accuracies into the frequency domain (after zero padding and application of a Hanning window). The FFT length was 160 data points with a Hanning window of 40 data points. A randomization procedure was performed to assess the statistical significance of the observed spectral amplitude. The ISI labeling of the detrended decoding accuracies was shuffled for each observer, and then FFT was performed on this surrogate data, similar to that of the original data analysis. This randomization procedure was repeated for 1000 times, generating a distribution of spectral amplitude for each frequency, from which a 0.05 (uncorrected) significance level was obtained.

### Phase difference analysis

To estimate the phase relationship between the left and right FFAs in face processing, using a similar FFT procedure in which the time-course of left and right FFA decoding accuracy was detrended, zero-padded, Hanning tapered, and then 160 data point Fourier transformed, we calculated and averaged the phase difference between the left and right FFA as a function of frequency from 0 to 15 Hz (30 Hz sampling rate) in each visual field condition for each observer. A further cross-observer coherence in the phase difference was calculated, and a left-right FFA phase difference coherence pattern was plotted as a function of frequency from 0 to 15 Hz. A similar randomization procedure as the frequency analysis was performed, and a 0.05 threshold (uncorrected) significance level was obtained. Multiple comparison correction was then applied to the uncorrected randomization threshold by setting the maximum across all frequency bins as the final permutation threshold of significance for cross-observer coherence in the left-right FFA phase difference. In each visual field condition (LVF-RVF and RVF-LVF), we tested the non-uniformity for left-right FFA phase differences across observers using circular statistics separately, in the significant frequency bands (Rayleigh test for circular data non-uniformity test in CircStats toolbox for MATLAB).

